# Independent pseudogenization of *CYP2J19* in penguins, owls and kiwis implicates gene in red carotenoid synthesis

**DOI:** 10.1101/130468

**Authors:** Christopher A. Emerling

## Abstract

Carotenoids have important roles in bird behavior, including pigmentation for sexual signaling and improving color vision via retinal oil droplets. Yellow carotenoids are diet-derived, but red carotenoids (ketocarotenoids) are typically synthesized from yellow precursors via a carotenoid ketolase. Recent research on passerines has provided evidence that a cytochrome p450 enzyme, CYP2J19, is responsible for this reaction, though it is unclear if this function is phylogenetically restricted. Here I provide evidence that CYP2J19 is the carotenoid ketolase common to Aves using the genomes of 65 birds and the retinal transcriptomes of 15 avian taxa. *CYP2J19* is functionally intact and robustly transcribed in all taxa except for several species adapted to foraging in dim light conditions. Two penguins, an owl and a kiwi show evidence of genetic lesions and relaxed selection in their genomic copy of *CYP2J19*, and six owls show evidence of marked reduction in *CYP2J19* retinal transcription compared to nine diurnal avian taxa. Notably, none of these taxa are known to use red carotenoids for sexual signaling and several species of owls and penguins represent the only birds known to completely lack red retinal oil droplets. The remaining avian taxa belong to groups known to possess red oil droplets, known or expected to deposit red carotenoids in skin and/or plumage, and/or frequently forage in bright light. The loss and reduced expression of *CYP2J19* is likely an adaptation to maximize retinal sensitivity, given that oil droplets reduce the amount of light available to the retina.

## 1. Introduction

Carotenoids perform numerous functions in animals (Olson and Owens, 1998), with much of the research on avian fauna focusing on its roles in sexual signaling (Gray, 1996; McGraw et al., 2002; Blount et al., 2003; Blas et al., 2006) and fine-tuning of color vision (Vorobyev et al., 1998; Vorobyev, 2003; Stavenga and Wilts, 2014; Toomey et al., 2015). Carotenoids can be deposited in highly visible areas of the body, including skin, plumage and beaks (Blount et al., 2003; Olson and Owens, 2005; Thomas et al., 2014) and are thought to provide honest signaling of health to potential mates (Gray, 1996; Hill and Johnson, 2012; Mundy et al., 2016). Carotenoids are also deposited in oil droplets located in retinal cone photoreceptors (Toomey et al., 2015), the cells that facilitate color vision. Here, they are believed to function as microlenses (Stavenga and Wilts, 2014) and act as long pass wavelength filters that minimize spectral sensitivity overlap between different cone classes (Vorobyev et al., 1998; Vorobyev, 2003), thereby enhancing color vision.

Though both yellow and red carotenoids are deposited on external anatomy and in retinal oil droplets of birds, these colors are typically produced via different pathways. Most birds obtain yellow carotenoids directly from their diets but can convert these into red ketocarotenoids via a ketolation pathway (Brush, 1990; Koch et al., 2016). Recent evidence has implicated a cytochrome P450, CYP2J19, in the endogenous synthesis of red carotenoids in birds. Lopes et al. (2016) examined the basis for red plumage in red factor canaries, a breed created by crossing normally yellow common canaries (*Serinus canaria*) with red siskins (*Spinus cucullata*) and repeatedly backcrossing these hybrids with common canaries. Multiple genome comparisons strongly suggest that the siskin red plumage locus was introgressed into the red factor canary genome, with two loci being strongly associated with the red plumage phenotype. Not only is *CYP2J19* found at one of these loci, Lopes et al. (2016) demonstrated that *CYP2J19* RNA expression is upregulated in the skin and livers of red canaries relative to yellow canaries, and is also expressed in growing red feathers. Finally, they demonstrated that both red and yellow canaries express *CYP2J19* in their retinas, with no significant differences between the two, suggesting that this gene is being used to make red retinal oil droplets. Mundy et al. (2016) examined another passerine bird, the zebra finch (*Taeniopygia guttata*), which sequesters ketocarotenoids in its beak and tarsi. However, a mutation called *yellowbeak* results in the absence of ketocarotenoids in both structures. Mundy et al. (2016) mapped the *yellowbeak* mutation to a 2.83 Mb region of the genome, which included two copies of *CYP2J19*. Notably, yellowbeak zebra finches are homozygous for a deletion of one of the copies, strongly implicating this locus. *CYP2J19* was further found to be expressed in the beak and tarsus of wild-type zebra finches and the retinas of both wild-type and *yellowbeak* finches, but is barely detectable in the beaks of *yellowbeak* individuals.

What is not clear is if *CYP2J19* is the same gene responsible for red carotenoids across avian taxa or is merely a passerine innovation. Twyman et al. (2016) found evidence that *CYP2J19* originated in the stem archelosaurian lineage (Testudinata + Archosauria), being present in the genomes of three testudines, and additionally found the gene transcribed in the retina and red portions of the plastron of western painted turtles (*Chrysemys picta*). This seems suggestive of a long history of red carotenoid production in archelosaurs using *CYP2J19*, though it could alternatively be explained by independent co-option of the same gene for carotenoid ketolase function. An additional means of testing this hypothesis is by examining the genomes of species that are likely to have lost the capacity to synthesize ketocarotenoids. If *CYP2J19* is indeed the avian carotenoid ketolase, then it should be deleted or pseudogenized in the genomes of species in which there is no selective pressure to maintain its function. Alternatively, if its importance has been diminished, it should be transcribed at substantially lower levels than species with abundant red carotenoid expression. A scenario that would lead to the loss of this gene should satisfy at least two conditions: (1) the species does not deposit red carotenoids externally for sexual signaling or other functions, and (2) the species does not sequester red carotenoids in oil droplets for color vision.

The first requirement for gene loss has been satisfied by many species. The phylogenetic distribution of species known and predicted to sequester ketocarotenoids in their skin and/or feathers implies that many lineages of birds have independently derived the ability to externally express these pigments (Olson and Owens, 2005; Thomas et al., 2014). Many species rely on different color mechanisms for sexual signaling (Stoddard and Prum, 2011), including structural coloration, porphyrins (turacos; Church, 1869; Rimington, 1939), spheniscins (penguins; McGraw et al., 2007; Thomas et al., 2013), psittacofulvins (parrots; McGraw and Nogare, 2004, 2005) or yellow and orange carotenoids. Other species have minimal coloration, having instead selected for more cryptic plumage, such as the nocturnal predatory owls and the marine-adapted penguins. By contrast, the loss of red carotenoids in retinal oil droplets is presumably much less frequent given that sharp color vision is presumably adaptive for most of the predominantly diurnal Aves. Indeed, red droplets appear to be very widespread, being found in nearly all birds queried across at least 22 orders of palaeognaths, Galloanseres and Neoaves (Krause, 1894; Erhard, 1924; Dücker, 1963; Strother, 1963; Peiponen, 1964; Mayr, 1972; Muntz, 1972; Yew et al., 1977; Sillman et al., 1981; Goldsmith et al., 1984; Jane and Bowmaker, 1988; Partridge, 1989; Bowmaker et al., 1993; Gondo and Ando, 1995; Bowmaker et al., 1997; Hart et al., 1998; Das et al., 1999; Hart et al., 2000a, b; Wright and Bowmaker, 2001; Hart, 2001; Coyle et al., 2012; Porter et al., 2014). However, carotenoid oil droplets are likely to be maladaptive under the conditions of dim light. Since their contribution to refining color vision involves light filtration, it may be disadvantageous for species adapted to dim light conditions to reduce the amount of light available to the retina. Indeed, the absence of carotenoids in retinal oil droplets, or oil droplets in general, in mammals, snakes, crocodylians and geckos has been used as evidence that these lineages have long histories of dim light adaptation (Walls, 1942; Röll, 2000). Among birds, red oil droplets are reported to be lacking or extremely rare in at least some penguins and owls (Erhard, 1924; Bowmaker and Martin, 1978; Bowmaker and Martin, 1985; Gondo and Ando, 1995), suggesting that minimizing carotenoid deposition may be an adaptation to aquatic and nocturnal predation.

I tested this hypothesis using publicly available genomes for 65 species of birds representing 36 taxonomic orders and 55 families and retinal transcriptomes from an additional 15 species. Here I describe evidence that the *CYP2J19* locus is inactivated in primarily dim light-adapted taxa and is only minimally transcribed in owl retinas, further supporting the hypothesis that *CYP2J19* is the locus responsible for red carotenoid-based coloration and color vision in birds.

## 2. Materials and methods

### 2.1. Assembling CYP2J19 sequences

I designed an *in silico* probe for obtaining genome-derived *CYP2J19* sequences by BLASTing (blastn) a *CYP2J19* gene model (chicken, *Gallus gallus*; XM_422553) against the chicken genome assembly in the NCBI Whole Genome Shotgun Contig database (WGS), and obtaining a contiguous sequences that included all exons, introns and some flanking sequence of the gene. I then BLASTed (discontiguous megablast) this probe against avian genomes in the WGS and aligned the resulting hits to the reference probe using MUSCLE (Edgar, 2004) in Geneious ver. 9.1.8 (Kearse et al., 2012; ESM, Dataset S1).

I obtained retinal transcriptome sequences by BLASTing (discontiguous megablast) the chicken *CYP2J19* gene model against 15 retinal transcriptomes deposited by Wu et al. (2016) in NCBI’s Sequence Read Archive database (SRA). I imported the BLAST hits for each species into Geneious, and assembled *CYP2J19* mRNAs by mapping the reads to the chicken reference using the Geneious assembler (medium sensitivity). For the owl species, the mapped BLAST hits frequently belonged to other cytochrome p450 gene paralogs and frequently assembled chimeric sequences, due to the low number of short reads. To ensure more accurate sequence assemblies, I also mapped these reads to the *Tyto alba CYP2J19* contig and manually removed portions of reads that appeared non-homologous. For each retinal transcriptome, I recorded the number of BLAST hits, reads mapped, mean coverage, maximum coverage, and percentage of the gene that was assembled. I compared the means of these metrics in owls versus the remaining diurnal taxa using Welch’s t-tests.

After aligning all *CYP2J19* sequences using MUSCLE in Geneious, I manually adjusted them (electronic supplementary material [ESM], Dataset S1, Table S1) and examined them for inactivating mutations, i.e., splice site mutations, frameshift insertions and deletions, and premature stop codons.

### 2.2. Molecular evolutionary analyses

I estimated the gene tree for avian *CYP2J19* from an alignment of genome-and retinal transcriptome-derived sequences and outgroup cytochrome P450 genes (ESM, Dataset S1). I performed a maximum likelihood analysis implementing RAxML ver. 8.2.9 (Stamatakis, 2014) in CIPRES (phylo.org; RAxML-HPC2 on XSEDE) using the default parameters and performing 500 bootstrap iterations.

I tested for evidence of relaxed selection on putative *CYP2J19* pseudogenes by performing dN/dS ratio (ω) analyses with codeml in PAML ver. 4.8 (Yang, 2007; ESM, Table S2). I omitted owl sequences derived from transcriptomes, removed stop codons and sites with dubious homology (ESM, Dataset S1), and assumed the topology from Prum et al. (2015). I performed analyses with both F1X4 and F3X4 codon frequency models, which differ in calculating frequencies from base composition versus base composition given codon position, respectively. All analyses involved branch models, which assume one ω for the entire gene and differing ω estimates for designated branches. I performed progressively nested models, with a single ω for the entire tree, followed by the estimation of additional individual branches expected to be evolving neutrally.

## 3. Results

### 3.1 CYP2J19 *gene tree estimation*

*CYP2J19* was reconstructed as monophyletic with 100% bootstrap support (ESM, Figure S1). Many well-established avian clades were also reconstructed as monophyletic, including Bucerotiformes, Piciformes, Accipitriformes, Falconiformes, Passeriformes, Psittaciformes, Apodiformes, Sphenisciformes, Mirandornithes, Charadriiformes, Palaeognathae, Galloanseres, Anseriformes and Galliformes. Strigiformes came out as paraphyletic with the strigiform *Otus bakkamoena* recovered with *Cuculus canorus* (Cuculiformes). However, the *O. bakkamoena CYP2J19* assembly only covered 10.2% of the total sequence, suggesting that this erroneous placement is a sampling artifact. Nearly all interordinal relationships conflict with the species tree, though this is likely due to a combination of limited sequence information and long branch attraction caused by non-optimal taxon sampling density (Prum et al., 2015).

### 3.2. CYP2J19 *pseudogenes*

Among the whole genome assemblies examined, I found evidence of genetic lesions in *CYP2J19* for seven species: the emperor penguin (*Aptenodytes forsteri*; Sphenisciformes), Adélie penguin (*Pygoscelis adeliae*; Sphenisciformes), barn owl (*Tyto alba*; Strigiformes), southern brown kiwi (*Apteryx australis*; Apterygiformes), red-throated loon (*Gavia stellata*; Gaviiformes), scarlet macaw (*Ara macao*; Psittaciformes), and MacQueen’s bustard (*Chlamydotis macqueeni*; Otidiiformes). For three of these species, the evidence for *CYP2J19* inactivation is relatively weak. *Ara macao* has a 1-bp deletion in exon 2 of *CYP2J19*, though this mutation was only found in a lower coverage assembly for the same individual (16× versus 26×), likely indicating an assembly error. *A. macao* did have an elevated dN/dS ratio (ω = 0.3466-0.3747), but this was not significantly different from the background ω (p = 0.064-0.066). The *Chlamydotis macqueeni* sequence has a 5-bp deletion at the intron 6 – exon 7 boundary, but BLASTing this region against the SRA and mapping the short reads back onto the intron – exon boundary indicates that this is most probably an assembly error. Consistent with this hypothesis, dN/dS ratio modeling found the *C. macqueeni* branch to have a dN/dS ratio (ω = 0.1226-0.1463) slightly lower than the background (ω = 0.1682-0.181), indicating purifying selection. In the loon, *Gavia stellata*, there is a single premature stop codon in the final exon (exon 9) of the gene, though the location of the putative mutation (AAG → TAG) is 5-bp from the end of the contig, meaning that it is plausibly an assembly error. dN/dS ratio models seem suggestive of a higher rate of protein evolution (ω = 0.2961-0.3225) relative to the background rate (ω = 0.1682-0.181), but a likelihood ratio test did not find this model to be a significantly better fit (p = 0.11-0.12).

By contrast, the penguin, owl and kiwi sequences show stronger evidence of inactivating mutations and subsequent relaxed selection in *CYP2J19*, consistent with loss of function (Figure 1). *Aptenodytes forsteri* has a single splice acceptor mutation in intron 7, resulting from an AG → AA substitution. The guanine is conserved in all other avian genomes examined, suggesting that the canonical splice acceptor (AG) is critical for functional retention. BLASTing against the SRA and mapping the short sequence reads back to the contig reveals that out of 79 reads that map to this locus, 54 (68.4%) have an A at this site versus 25 (31.6%) that have a G. This implies that this individual is heterozygous at this site, and therefore AA may not be fixed in this species. Nonetheless, dN/dS ratio analyses reveal that selection is probably relaxed in *A. forsteri*, with an ω of 0.5101–0.5397 (p < 0.01). More convincingly, *Pygoscelis adeliae* has both a premature stop codon in exon 2 and a 1-bp deletion in exon 7, along with a very high dN/dS ratio of 0.8776–0.921 (p < 0.001). dN/dS ratios for the stem sphenisciform branch were low (ω = 0.2652–0.2869) and not significantly different from the background (ω = 0.1682-0.1798; p = 0.15-0.16), suggesting that selection was relaxed in parallel on the individual penguin descendant branches. *Tyto alba* has the largest number of loss-of-function mutations, including premature stop codons in exons 1, 5 and 6, and a 5-bp deletion in exon 3, as well as an ω of 0.6883–0.7392 (p < 0.001), indicative of relaxed selection on this branch. Finally, the nocturnal *Apteryx australis* has three genetic lesions, a 5-bp insertion in exon 5, an 8-bp deletion in exon 9, and exon 7 is completely deleted. Consistent with the prediction of gene inactivation, the *Apteryx australis* branch is estimated to have a high dN/dS ratio (ω = 0.8293–0.8358; p < 0.001).

**Figure 1.**
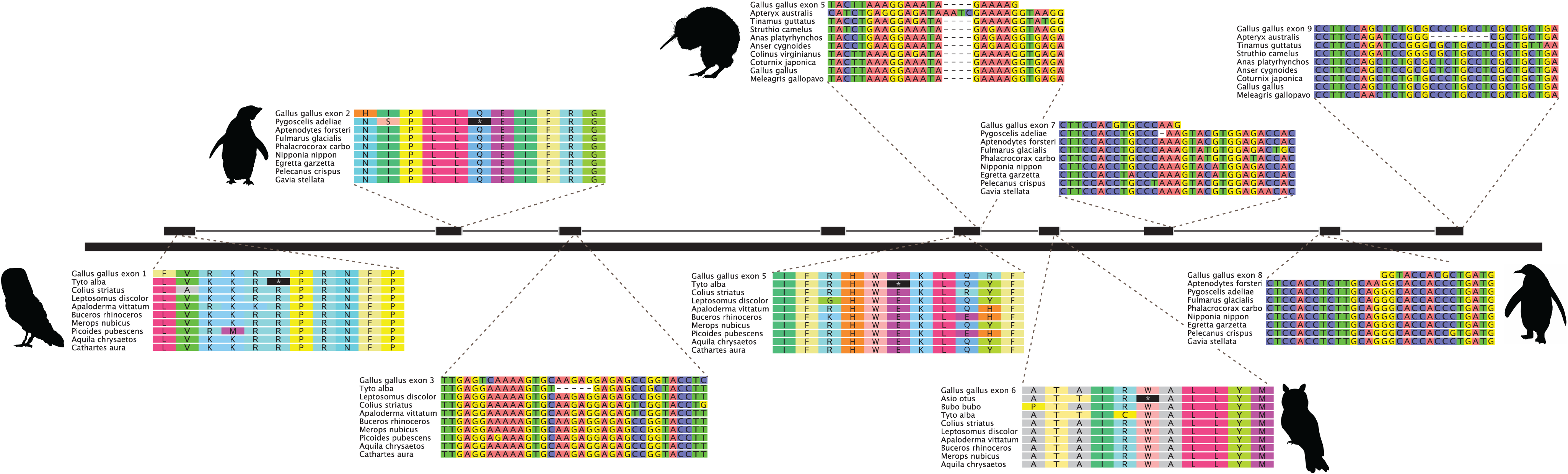
Examples of *CYP2J19* loss-of-function mutations in the barn owl (*Tyto alba*), long-eared owl (*Asio otus*), Adélie penguin (*Pygoscelis adeliae*), emperor penguin (*Aptenodytes forsteri*) and Southern brown kiwi (*Apteryx australis*). Examples include premature stop codons (black asterisk symbols in amino acid alignments), frameshift indels and a splice acceptor mutation (AA). Silhouettes from phylopic.org (see website for individual licenses).

### 3.3. CYP2J19 *retinal transcription comparisons*

I also tested for the presence of *CYP2J19* in published avian retinal transcriptomes (Wu et al. 2016), which encompass numerous diurnal avian taxa (Accipitriformes [five spp.], Falconiformes [two spp.], Bucerotiformes [hoopoe; *Upupa epops*], Piciformes [grey-headed woodpecker; *Picus canus*]) and six species of owls (Table 1). By mapping the reads to the *Gallus gallus* reference sequence, I was able to assemble a complete or nearly complete (99.9%) *CYP2J19* gene in all nine diurnal birds, with generally strong coverage in each species. By contrast, I was unable to assemble a complete *CYP2J19* transcript in any of the owls (23.1–71%), with much lower mean values for the total number of BLAST hits, reads mapped, average coverage and maximum coverage compared to the diurnal species, each difference being statistically significant (Table 1). The most complete strigiform transcript, which had the highest assembly metrics (*Asio otus*; long-eared owl), also showed evidence of being a transcribed pseudogene (Figure 1), possessing a 4-bp deletion in exon 1 (2× coverage) and premature stop codons in exons 6 (8× coverage) and 8 (5× coverage).

**Table 1.** Metrics of *CYP2J19* assemblies derived from retinal transcriptomes.

| Order | Species | BLAST hits | Reads mapped | Mean coverage | Maximum coverage | % of ref seq |
| --- | --- | --- | --- | --- | --- | --- |
| **Strigiformes** | *Asio otus* | 261 | 105 | 8.6 | 67 | 71% |
|  | *Athene noctua* | 175 | 67 | 5.7 | 38 | 45.70% |
|  | *Bubo bubo* | 144 | 58 | 4.9 | 36 | 38.30% |
|  | *Otus bakkamoena* | 22 | 7 | 0.5 | 5 | 23.10% |
|  | *Otus scops* | 138 | 68 | 5.6 | 42 | 64.90% |
|  | *Tyto longimembris* | 28 | 18 | 1.5 | 9 | 31.10% |
|  | **Strigiformes** | **128** | **53.8** | **4.5** | **32.8** | **45.70** |
| **Accipitriforme** | *Accipiter virgatus* | 3453 | 3135 | 258.9 | 429 | 100% |
|  | *Aegypius* | 3546 | 3370 | 277.1 | 461 | 100% |
|  | *Bustatur indicus* | 2805 | 2495 | 207.1 | 343 | 100% |
|  | *Circus* | 1206 | 1086 | 90.4 | 162 | 100% |
|  | *Elanus caeruleus* | 262 | 209 | 17.5 | 36 | 100% |
| **Falconiformes** | *Falco subbuteo* | 6700 | 6274 | 517.9 | 1057 | 100% |
|  | *Falco tinnunculus* | 6274 | 6300 | 525.9 | 925 | 100% |
| **Piciformes** | *Picus canus* | 2334 | 1790 | 146.6 | 403 | 99.90% |
| **Bucerotiform** | *Upupa epops* | 2254 | 2026 | 168.7 | 305 | 100% |
|  | **Non-Strigiformes Mean** | **3204** | **2965** | **245.6** | **457.9** | **100%** |
|  | **Welch's t-test p-values** | **0.002** | **0.003** | **0.003** | **0.005** | **0.0009** |

## 4. Discussion

### 4.1. Inactivation of CYP2J19 in dim light-adapted birds

Here I provide molecular evolutionary evidence that *CYP2J19* is the avian carotenoid ketolase locus, corroborating previous studies of passerines (Lopes et al., 2016; Mundy et al., 2016). Out of 65 avian genomes and 15 retinal transcriptomes I examined, encompassing 57 families of birds, *CYP2J19* is intact and/or transcribed robustly in all but a few species. These exceptions include nocturnal (owls, kiwi) and aquatic predators (penguins), and encompass the only two clades (Strigiformes, Sphenisciformes) that have species known to lack both external and retinal red carotenoids.

The Humboldt (*Spheniscus humboldti*) and king penguins (*Aptenodytes patagonicus*) both lack red retinal oil droplets, with the former possessing yellow, pale and clear oil droplets (Bowmaker and Martin, 1985) and the latter only having pale-green droplets (Gondo and Ando, 1995). By contrast, the rockhopper penguin (*Eudyptes chrysocome*) was reported to have four colors of oil droplets (Gondo and Ando, 1995), more typical of diurnal birds, though the specific colors were not reported. Regardless, the fact that there were no shared genetic lesions between *Aptenodytes forsteri* and *Pygoscelis adeliae*, and the dN/dS ratio estimates do not indicate complete relaxation of selection on their respective branches, suggests that penguins lost red oil droplets in parallel, rather than on the stem Sphenisciformes branch. The retention versus loss of these droplets in specific penguins may be due to differences in light exposure among the various taxa. Notably, Adélie and emperor penguins, both of which have genetic lesions in *CYP2J19*, are among the deepest diving penguins (Schreer and Kovacs, 1997) and have an exclusively circum-Antarctic distribution. Diving in deeper water and exposure to extensive periods of darkness during the winter may have been important selection pressures leading to *CYP2J19* loss.

Some owls are reported to possess red oil droplets, including the short-eared (*Asio flammeus*), little (*Athene noctua*), and tawny owls (*Strix aluco*), whereas others, such as the snow (*Bubo scandiacus*), Ural (*Strix uralensis*) and barn owls (*Tyto alba*) appear to lack them (Erhard, 1924; Yew et al., 1977; Bowmaker and Martin, 1978; Gondo and Ando, 1995). Even in the tawny owl, red pigments are limited to less than 1% of all oil droplets (Bowmaker and Martin, 1978), suggesting that even when retained in owls, they are less abundant. In addition to the evidence of *CYP2J19* pseudogenization in the barn owl (Tytonidae), Hanna et al. (under review) reported that the spotted owl’s (*Strix occidentalis*) copy of *CYP2J19* has a single frameshift mutation and also shows evidence of being under relaxed selection. These data, along with the putative transcribed *CYP2J19* pseudogene in the long-eared owl and overall minimal expression of *CYP2J19* in owls compared to other avian species, are consistent with a diminished importance of red carotenoid oil droplets in these nocturnal predators.

The retinal oil droplets of the nocturnal kiwis have not been characterized, but the fact that these birds are adapted to dim light, with retinal histology highly similar to barn owls (Corfield et al., 2015), suggests that they are likely to lack red oil droplets in their retinas. The presence of a *CYP2J19* pseudogene and strong evidence for relaxed selection on this gene further supports this hypothesis, and should be evaluated in future studies.

An additional example of *CYP2J19*’s link to red carotenoid oil droplets comes from the genomes of crocodylians. Twyman et al. (2016) found evidence that *CYP2J19* arose in the stem archelosaurian lineage due to its presence in some turtles and birds and absence in other vertebrates. However, Twyman et al. (2016) could not find the gene in the genomes of both *Alligator* species, and I was unable to find *CYP2J19* in the Nile crocodile (*Crocodylus porosus*) or Indian gharial genomes (*Gavialis gangeticus*), potentially indicating whole gene deletion in the crocodylian ancestor. Notably, crocodylians completely lack retinal oil droplets, which are thought to have been lost in their last common ancestor during a ‘nocturnal bottleneck’ along with other traits associated with light detection (Walls, 1942; Emerling, 2017). Together, the loss of *CYP2J19* across dim light-adapted birds and crocodylians supports Twyman et al.’s (2016) hypothesis that it is a uniquely archelosaurian carotenoid ketolase.

### 4.2 Retention of CYP2J19 across most avian taxa

Most avian taxa retain an intact *CYP2J19* gene, suggesting retention of the ability to synthesize ketocarotenoids, a conclusion consistent with the widespread retention of red cone oil droplets. Some of these species forage under dim light conditions, including the nocturnal/crepuscular Chuck-will’s-widow (*Caprimulgus carolinensis*; Caprimulgiformes) and chimney swift (*Chaetura pelagica*; Apodiformes) and the diving great cormorant (*Phalacrocorax carbo*), great crested grebe (*Podiceps cristatus*) and red-throated loon (*Gavia stellata*). The related European nightjar (*Caprimulgus europaeus*) and common swift (*Apus apus*) are reported to retain red oil droplets, as are some diving ducks (Anseriformes) and cormorants (Suliformes) (Krause, 1894; Muntz, 1972; Hart, 2001). However, these droplets are relatively depauperate in such species. For example, Hart (2001) found only 6.7% of oil droplets in the pied cormorant retinal are red, much lower than in 21 other birds he examined (range = 12.5–35.6%; mean = 20.2%). Similarly, in the European nightjar and common swift, red droplets make up <10% and <6% of the total number of oil droplets, respectively (Krause, 1894; Muntz, 1972). These data suggest that although reducing the amount of red oil droplets may be common in dim light-adapted species, their complete loss is relatively rare, and may be indicative that penguins, owls and kiwis are among the best adapted to dim light. Indeed, among avian taxa, only owls (Bowmaker and Martin, 1978; Borges et al., 2015; Wu et al., 2016; Hanna et al., under review), penguins (Bowmaker and Martin, 1985; Li et al., 2014; Borges et al., 2015) and kiwis (Le Duc et al., 2015) show evidence of reduction in the number of cone visual pigment classes, an otherwise common pattern among vertebrates adapted to dim light (Douglas et al., 1995; Meredith et al., 2013; Jacobs, 2013; Emerling and Springer, 2014; Emerling, 2017).

There are a number of species with an intact *CYP2J19* gene belonging to orders with no available evidence of red retinal oil droplets. Two of these species hail from orders that are confirmed to deposit red carotenoids in the skin and plumage: the white-tailed tropicbird (*Phaethon lepturus*; Phaethontiformes) and the bar-tailed trogon (*Apaloderma vittatum*; Trogoniformes; Olson and Owens, 2005; Thomas et al., 2014). Several other species come from clades with skin coloration consistent with red carotenoids: the rhinoceros hornbill (*Buceros rhinoceros*; Bucerotiformes), red-legged seriema (*Cariama cristata*; Cariamiformes), speckled mousebird (*Colius striatus*; Coliiformes), cuckoo roller (*Leptosomus discolor*; Leptosomiformes), MacQueen’s bustard (*Chlamydotis macqueeni*; Otidiformes), and yellow-throated sandgrouse (*Pterocles gutturalis*; Pteroclidiformes; Olson and Owens, 2005). The retention of *CYP2J19* in each of these species highlights the probability that their red skin coloration is indeed derived from ketocarotenoids, though this will need to be validated. Finally, several species come from orders that have neither been examined for red oil droplets nor appear to have plumage or skin pigmentation derived from red carotenoids: the sunbittern (*Eurypyga helias*; Eurypygiformes), red-throated loon (*Gavia stellata*; Gaviiformes), brown mesite (*Mesitornis unicolor*; Mesitornithiformes), red-crested turaco (*Tauraco erythrolophus*; Musophagiformes), hoatzin (*Opisthocomus hoazin*; Opisthocomiformes), and great crested grebe (*Podiceps cristatus*; Podicipediformes; Olson and Owens, 2005; Thomas et al., 2014). Despite the apparent absence of red carotenoid pigmentation, the retention of *CYP2J19* suggests that this gene is at least being maintained for the synthesis of red retinal oil droplets.

The retention of functional *CYP2J19* orthologs across diverse avian taxa raises questions regarding alternative mechanisms for producing red pigmentation. For instance, parrots produce the clade-specific red psittacofulvins despite the presence of circulating serum carotenoids (McGraw and Nogare, 2004), red oil droplets in the retinas of cacatuid and psittaculid parrots (Bowmaker et al., 1997; Hart, 2001) and an intact *CYP2J19* in psittacid (blue-fronted amazon, *Amazona aestiva*; *Ara macao*), psittaculid (budgerigar, *Melopsittacus undulatus*) and strigopid parrots (kea, *Nestor notabilis*). Why did parrots evolve a novel pigment rather than co-opting *CYP2J19*, similarly to what many other birds appear to have done? Additionally, can the ability to synthesize psittacofulvins for external coloration lead to co-option of this pigment for color vision in place of red carotenoids? Turacos also have a unique red pigment, known as turacin (Church, 1869; Rimington, 1939), despite the fact that at least one species (*Tauraco erythrolophus*) has an intact *CYP2J19* gene. Future research will need to confirm the presence of red oil droplets in the retinas of turacos, but for now it raises the question as to why they evolved novel red pigments when the genetic machinery for producing red carotenoids was present in their genomes.

## 5. Conclusion

Here I provided molecular evolutionary evidence that *CYP2J19* is the carotenoid ketolase common to Aves, bolstering evidence from recent research in passerines and the painted turtle (Lopes et al., 2016; Mundy et al., 2016; Twyman et al., 2016). The pseudogenization and relaxed selective constraints on this gene in species adapted to foraging in dim light niches, and belonging to clades known or expected to lack red carotenoid oil droplets, plumage and skin pigmentation, provides strong evidence that this gene is tied to the synthesis of ketocarotenoids. Additionally, retinal transcriptomes in nocturnal owls showed a marked decrease in transcription of this gene compared to diurnal outgroup taxa, consistent with the absence or reduction in red retinal oil droplets in owls. The pattern of *CYP2J19* loss/reduction in species adapted to dim light highlights the likelihood that this is an adaptation to minimize light filtration, thereby enhancing visual sensitivity in darkness.

## Acknowledgments

This research was supported by an NSF Postdoctoral Research Fellowship in Biology (Award #1523943).

